# Quantum Convolutional HLA Immunogenic Peptide Prediction (Q-CHIPP): Next-Generation Neoantigen Prediction with Quantum Neural Networks

**DOI:** 10.1101/2025.07.29.667313

**Authors:** Ryan Peters, Kahn Rhrissorrakrai, Prerana Bangalore Parthasarathy, Vadim Ratner, Tanvi P. Gujarati, Meltem Tolunay, Jie Shi, Jeffrey K. Weber, Timothy A. Chan, Laxmi Parida, Sara Capponi, Filippo Utro, Tyler J. Alban

## Abstract

The immune system is an intricately evolved series of cellular and protein-protein interactions, which defend the body against pathogens and abnormal cells such as cancer. A key player in the immunologic recognition of non-self is the immune synapse, where T cell receptors (TCRs) scan peptides presented on major histocompatibility complex (MHC) molecules to detect and eliminate cells displaying non-self antigens. While this interaction is vital for vaccine and immunotherapy success, the underlying rules of TCR recognition remain poorly understood. This is only further challenged as the application of predictive models is very limited due to small training datasets. While traditional machine learning models excel at predicting neoantigen binding to MHC, they often struggle to accurately predict immunogenicity. To address these challenges, we developed a quantum computing approach using Quantum Convolutional Neural Networks (QCNNs). Here we present Quantum Convolutional Human Leukocyte Antigen (HLA) Immunogenic Peptide Prediction (Q-CHIPP), the first application of quantum hardware based on training/predicting both MHC binding and immunogenicity in a combinatorial approach. Additionally, we present a large scale use of quantum hardware at scale with 46 qubits. This study underscores how quantum technology can be used for biological modeling and presents a scalable QCNN design with the potential to overcome current computational bottlenecks as quantum hardware advances.

## Introduction

Advancements in computational modeling have changed the way we approach scientific questions; however, many niche areas still face computational challenges due to limited training data, often within highly complex systems^1^. One problem instance plagued by these issues is the ability to accurately predict the targets of CD8 + T cells^2^. Predicting immunogenic epitopes that are targets of CD8+ T cells has become increasingly critical for the development of vaccines and cancer immunotherapy^3,4^. These immunogenic peptides exhibit characteristics linked to both peptide presentation and TCR recognition, with attributes related to MHC presentation often having a more pronounced influence due to conserved anchor positions and specific motifs associated with different HLA types^5^. Specifically, anchor residues at positions 1, 2 and 9 are known to strongly influence MHC binding, while TCR contact residues at positions 4 and 5 are involved in TCR contact (Fig. 1A). We recently explored these interactions in depth through molecular dynamic simulations of HLA binding neoantigens^6^ and showed that complex time-dependent events can be missed by oversimplifying interactions to fit current computational hardware.

**Figure 1.**
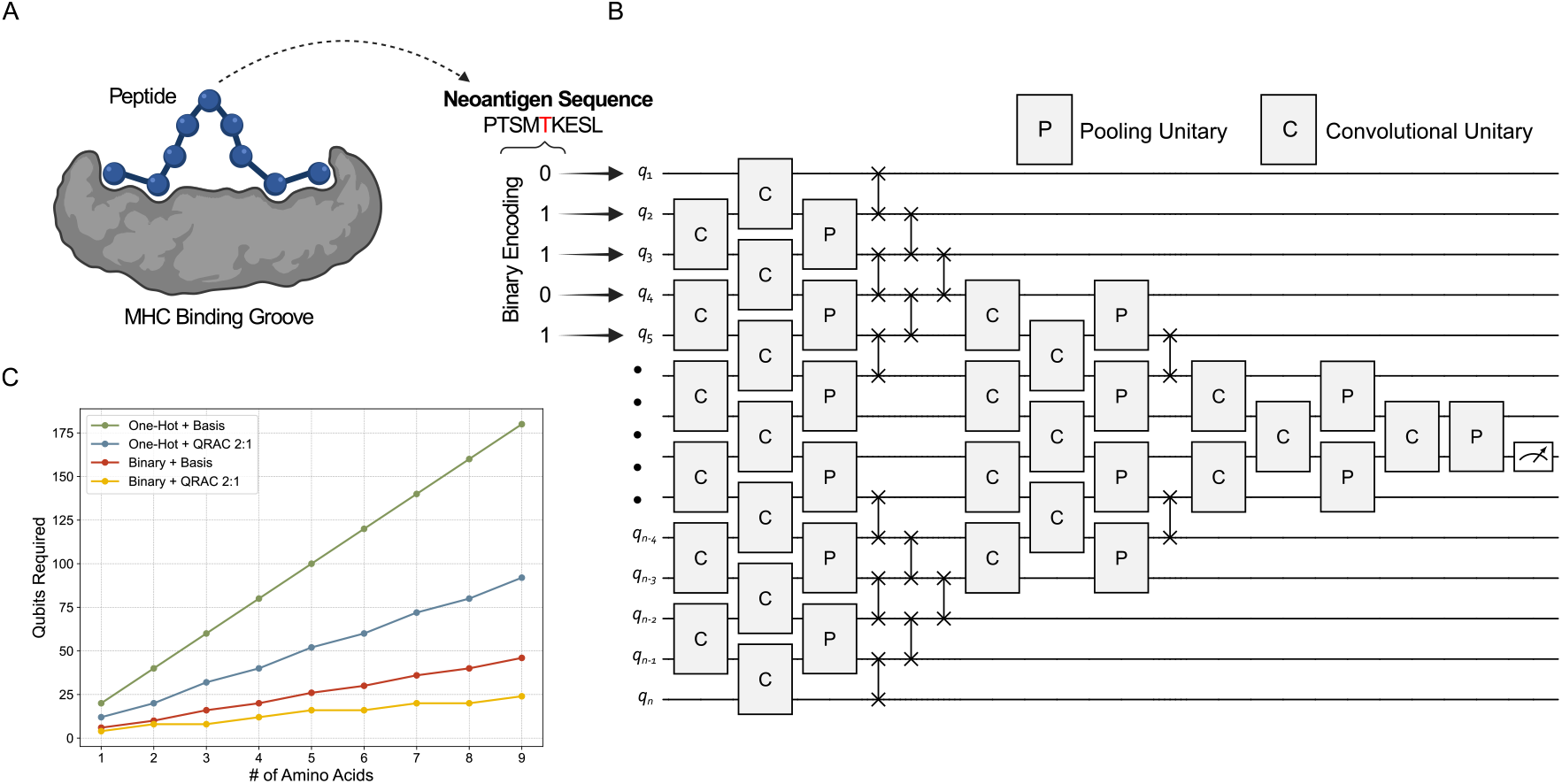
Encoding and quantum circuit design for MHC binding classification. A) Diagram of 9-mer neoantigen peptide binding in the MHC groove. Each amino acid is first converted to a 5-bit binary vector and is then mapped to the quantum circuit through the selected feature map (basis or QRAC 2:1). B) Schematic of the Quantum Convolutional Neural Network circuit. The circuit is composed of a convolutional layer followed by pooling layer. Each convolutional layer consist of two-qubit convolutional unitaries (C) on alternating nearest neighboring pairs of qubits. This is followed by a pooling layer composed of the two-qubit pooling unitary (P) which consists of 2 CNOT gates and 3 parameters. These layers are followed by a series of swap gates in order to uniformly discard the top and bottom set of qubits (pyramid circuit). C) Number of qubits required for different encoding schemes (one-hot & binary) and feature maps (basis & QRAC 2:1) across varying number of amino acids. Note that the QRAC 2:1 mapping requires an even number of classical bits to map correctly to the quantum space. Additionally, the final number of qubits, after all encodings and mappings, must be even to be compatible with the pyramid circuit architecture. As a result, classical data may be padded as by 1-3 extra bits to meet these constraints.

In addition to the complexity of MHC/peptide/TCR interactions, experimental limitations complicate the generation of training data because of the intrinsic lack of definitive true negatives. This is because a negative T cell assay indicates that a peptide failed to induce a T cell response in that specific experiment, but it does not necessarily mean that such peptide is empirically non-immunogenic. This issue was highlighted in a recent benchmarking exercise that compared all of the top artificial intelligence (AI) models (PRIME, DeepImmuno, iPred, NetTepi, REpitope, GAO, NetMHCpan) for predicting immunogenicity of SARS-CoV-2 antigens^2^. Briefly, these data demonstrated poor prediction accuracy (AUC=0.497-0.574) when tasked to predict immunogenic peptides from a novel antigen^2^. Training data is further limited by high experimental costs and the patient/HLA specific nature of screening that leads to biases based on currently available tools. Another study highlighting the extent of this problem was carried out by a consortium of 28 teams in academia and industry, where they screened 608 peptides and ultimately identified 37 immunogenic peptides (hit rate 6% based on model prediction methods in the field)^7^. Other groups have attempted to overcome this issue by pulling available data from multiple studies to train a type of meta-analysis model. This was the approach of the PRIME model, where data was collected from all available sources researchers could access and 129 true positive cancer neoantigens and 3200 negatives were identified^8^. While the model was one of the most advanced data mining efforts using state of the art machine learning (ML), it did not generalize well showing poor accuracy in predicting immunogenicity in a new dataset^2^. Recently we sought to rectify the lack of immunogenicity training data by developing the largest dynamic cancer immunogenicity screen to date^9^. This screen was unique in that it used paired samples prior to immune checkpoint therapy and post-therapy. This helped us enrich for immunogenic peptides by selecting those predicted antigens which were reduced on-therapy. With this method our success rate grew to 13%, roughly double the standard in the field.

In the last decade, interest in applying quantum computing (QC) in medicine-related fields has increased considerably^10–12^. QC leverages features of quantum mechanics, e.g. quantum entanglement and superposition, and could offer advantages over classical computing for certain computational problems. Quantum Machine Learning (QML) approaches may represent a potential solution for challenges observed in classical ML models^13,14^. Quantum algorithms could identify patterns and behavior present in classical or quantum data that may not be captured by classical algorithms or could identify solutions with fewer resources than those employed by classical algorithms^13,15^. Specifically, it has been demonstrated that Quantum Convolutional Neural Networks (QCNNs) generalize well with fewer training samples^16^, have better performances compared to their corresponding classical ML approaches^17,18^, and may offer some advantage over classical models when analyzing data^16^. On pre-fault tolerant quantum devices (PFTQDs), where noise is a critical factor, QCNN and other variational quantum algorithms must overcome challenges such as barren plateaus^19^, state preparation^20^, and advancements in classical computing in the form of quantum-inspired algorithms^21,22^. However, there remains an opportunity for the empirical demonstration of improved performance or novel biological insights as compared to state-of-the-art classical approaches using quantum computation.

In this study we applied our database of internal neoantigens and publicly available neoantigens to train and test QCNN models for both MHC binding prediction and immunogenicity prediction. Quantum experiments were executed on the IBM System One quantum processing unit (QPU) at The Cleveland Clinic and often found comparable performance despite relatively small circuit sizes and using a PFTQD. A QCNN approach may offer an advantage with learning with these limited numbers of samples, and provides certain scaling advantages over conventional quantum neural networks in the number of trained parameters important for testing on currently available quantum devices. Part of these investigations includes building the infrastructure to use QCNN for neoantigen prediction, testing noise mitigation techniques, and working to scale methods to full length neoantigens. While there are still limitations to current hardware, we ascertain the current feasibility of using these methods and develop a framework to expand with the ever-improving quantum hardware landscape. When testing the model on MHC-binding, the QCNN yielded F1 scores of 0.93 and 0.73 when using three or four amino acid positions, respectively, despite low training size of n=150 samples. We further demonstrate methods in modeling peptide strings with QCNN for immunogenicity prediction, showing ways to use a warm start method to optimize error mitigation and run-time. We also explore multiple encoding techniques, benchmarking to classical methods, and full length peptide analysis pushing the limits of current hardware with our largest model, successfully using 46 qubits. Additionally we showed that our QCNN model can be used to predict immunogenic antigen load which corresponds to immunotherapy response in a cohort of patients with lung cancer treated with immunotherapy. While existing ML/AI models have enhanced our understanding of T cell recognition and the development of vaccines and immunotherapies, we present here an exploration of how quantum computing may address their performance gaps to improve the identification of T cell targets in diverse immunological settings.

## Results

### QCNN encoding for MHC binding prediction

To enable immunogenic neoantigen prediction using quantum, we first focus on the development and benchmarking of the QCNN architecture for predicting peptide-MHC binding. In search of an appropriate quantum algorithm for neoantigen modeling, we selected QCNNs because of the empirical evidence of their suitability in data-limited problems^16^, in addition to the ubiquity of CNN-based tools in the field that offer a way to model antigens as peptide strings. As a foundational step, our approach centers on designing and evaluating data encoding strategies and circuit architectures using both noise-free simulators and quantum hardware. Here, we refine the quantum computing results using error mitigation techniques to ensure reliable performance on quantum hardware.

The QCNN circuit structure is composed of multiple blocks placed one after the other, such that at the end of one block half of the active number of qubits in that block remain. A schematic representation of the QCNN circuit is presented in Fig. 1B. Each block is composed of a convolution layer and a pooling layer. The convolution layer is implemented as a sequence of two-qubit unitaries called the convolution unitary (C) on alternating nearest neighboring pairs of qubits as shown. We tested the QCNN model with a convolutional unitary circuit, where 2-qubit gates are composed of 3 CNOT gates and 3 parametric single-qubit gates as a subspace of the universal KAK decomposition^23^. The convolution layer is then followed by the pooling layer. The pooling layer is composed of two-qubit pooling unitary (P) that acts on pairs of qubits (*q*_*i*_, *q*_*i*_ + 1) where *i* starts with qubit 1 or 2 based on whether total number of active qubits are a multiple of 4 or not for that layer. As shown in Fig. 1B the P blocks cover the full length of active qubits in each layer. Each pooling unitary has 2 CNOT gates and 3 free parameters. At the end of each pooling layer, we discard qubits symmetrically from the top and bottom of the register, retaining the 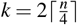 qubits where *n* is the number of qubits before pooling. To prepare for the next convolutional layer, we use a sequence of SWAP gates between adjacent qubits to align the central qubits into neighboring positions. The blocks performing the convolution and pooling operators are then repeated until only a single active qubit remains in the circuit. This qubit is then measured in the Z basis.

A crucial step in using QML algorithms to solve classical problems is the encoding of classical information into quantum states. The dataset used for this study is formed by MHC class I neoantigens of 9 amino acids in length, which considering all combinations of amino acids results in 512 million possible peptides that can bind more than 25,000 unique MHC I alleles. To limit this problem’s scope, we focus here on the most common HLA allele HLA-A*02:01 and begin by modeling only specific residues known to influence binding or T cell recognition. We investigate two encoding strategies for representing the peptide sequences. The first is one-hot encoding, in which each amino acid position is represented as a binary vector of length equal to the number of distinct amino acid classes observed. The second is binary encoding, which results in more compact representations by converting each amino acid to its base-2 equivalent, yielding a fixed-length 5-bit binary vector.

Unlike one-hot encoding, this method allows vectors to contain multiple non-zero entries, reducing matrix sparsity. In order to map the encoded classical data into each of the respective qubits, we utilize Z, ZZ, basis, and Quantum Random Access Coding (QRAC^24^) feature maps. Z or ZZ feature maps introduce one or two qubit unitary matrices, respectively, at the beginning of the QCNN circuit model to facilitate encoding the classical data into the quantum circuit via angles of Z rotation gates^15^. The basis feature map directly encodes binary input strings into quantum basis states by applying X gates to qubits corresponding to classical bits with value 1. Additionally, QRAC is utilized as a feature map compression scheme where 2 or 3 classical bits are encoded to a single qubit. Fig. 1C reports the number of qubits needed to map different amino acid sequence lengths using the encoding methods with each of the feature maps. Reducing the number of qubits is an important factor as it impacts the overall circuit size and determines what can be executable on current quantum hardware given levels of error and error mitigation.

To account for runtime efficiency and hardware constraints, we were required to limit the number of samples used in our training and testing splits. Our primary dataset, sourced from the Immune Epitope Database (IEDB), consisted of 8,740 total samples, of which 34% were classified as true positive MHC binders. Within these binders, 36% were further confirmed as immunogenic peptides. Note that on the quantum hardware, each circuit execution (i.e. forward pass) must be repeated multiple times, often referred to as shots, to gather multiple measurement outcomes to obtain expectation value of the measured observable. In addition, the noise present in current quantum hardware increases the need to measure more shots in order to mitigate the errors incurred due to noise. To balance the time taken to run the quantum experiments and get good measurement estimates, we were tasked with finding a balance between the number of shots and that of samples within each split. To address this, we curated a subset of the IEDB dataset consisting of 150 training samples and 50 test samples, each with a balanced distribution of positive and negative classes.

### Modeling MHC binding in quantum simulations

To begin modeling neoantigens with a quantum model, we started with a series of experiments varying the number and combinations of amino acid positions, aiming to evaluate the performance on a noiseless statevector simulator prior to migrating our solution to a PFTQD. We first focused on predicting MHC binding as there is greater data availability and the ground truth is more clear for binding assay based data than for immunogenicity. Our initial experiment consisted in evaluating the QCNN with three amino acid positions (positions 2, 5, and 9) along with NetMHCpan’s binding affinity rank (BA RANK). Here we chose to include BA rank when evaluating early models as an internal control parameter to help evaluate noise and encoding methods, understanding that BA rank has high accuracy in predicting binding. In order to minimize qubit requirements for computational efficiency on simulator, we applied binary encoding with a QRAC 2:1 feature map, resulting in an eight qubit experiment. After training, the model achieved an F1 score of 0.93 when tested upon the unseen dataset for MHC binding prediction, as seen in Table 1.

**Table 1.** Performance (F1 scores) for QCNN and CCNN models across input configurations. Classical models include both the limited parameter baseline to model the low number of parameters in a QCNN as well as the unrestricted baseline.

| Positions | QCNN | CNN (ltd.) | CNN (large) | RF |
| --- | --- | --- | --- | --- |
| BA | 0.89 | **0.90** | **0.90** | 0.88 |
| 2,5,9,BA | **0.93** | 0.84 | 0.90 | 0.88 |
| 2,4,5,9 | 0.80 | 0.66 | **0.88** | 0.83 |

Training involved stochastic parameter updates, with cross-entropy loss computed after each iteration to quantify prediction error. By tracking the loss over each iteration, we were able to determine whether the model was learning effectively. A consistent downward trend in the loss values indicates convergence towards an optimal set of parameters. To ensure proper training and establish that the model is converging, we analyzed the loss trajectories. These plots additionally helped us determine effective hyper-parameters, such as the number of iterations for training, learning rates, and effective error mitigation techniques, such as number of shots and usage of Pauli Twirling^25^ or Dynamical Decoupling^26–28^. By analyzing the loss plot for the initial three positions with BA RANK experiment, as seen in Fig. 2A, we see that around ~300 iterations, the loss values begin to converge after a sharp initial decrease. This indicates that the model had enough time (number of iterations) to train with effective learning rates. The high performance observed within this initial setup served as a positive control to the efficacy of our QCNN model since the performance was comparable with classical ML models (Table 1), which is expected given the integration of BA RANK along with peptide positions containing known MHC anchor residues.

**Figure 2.**
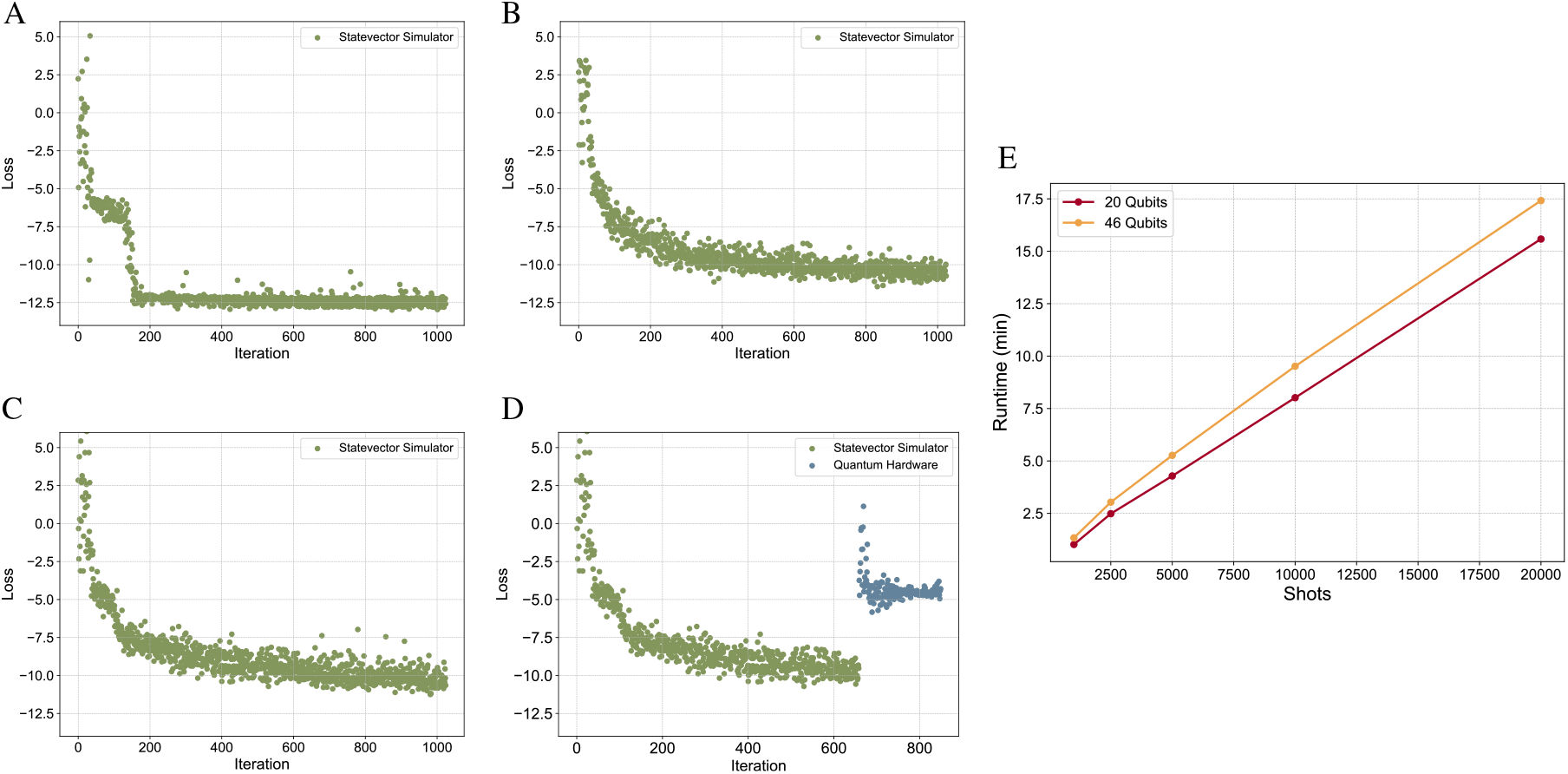
Impact of input configuration, feature map selection, and backend execution on QCNN training for MHC binding classification. Training loss curves for QCNN models under different input configurations and feature maps for initial MHC binding classification. Note all models used identical data splits (150 training and 50 test samples). A) Three peptide positions (2, 5, 9) in addition to BA RANK trained using noiseless statevector simulator with QRAC 2:1 feature map (8 qubits). B) Four peptide positions (2, 4, 5, 9) using a noiseless statevector simulator with QRAC 2:1 feature map (10 qubits). C) Same input features as (B) but using basis feature map on noiseless statevector simulator (20 qubits). D) Same input features and feature map as (C); model was initially trained on the noiseless simulator and then warm-started on quantum hardware (20 qubits). E) Runtimes on PFTQDs for 20 qubit experiment and 46 qubit experiment using different total number of shots.

After establishing the robustness of our model, we removed BA RANK and increased the number of encoded positions to four. This included binding residues (positions 2 and 9) and TCR recognition sites (positions 4 and 5). Implementing this model on the noiseless simulator resulted in an F1 score of 0.73 when mapped to the quantum states using QRAC 2:1 (10 qubit experiment). However, due to the data compression within this mapping and the potential for information loss, we additionally tested a 1:1 mapping to evaluate the extent to which the compression affects the model. To achieve this we employed a basis feature map which yielded a larger 20 qubit model. This resulted in an increased F1 score of 0.80 (Table 1). This is a particularly interesting result, given the small training size of *n* = 150 samples. We once again monitored the respective loss plots (Fig. 2B and 2C) to determine proper convergence and learning rates for our initial models.

### Warm start QCNN on QPU to predict MHC binding

Following our promising results on quantum simulators, we transferred our model to the quantum hardware and used the IBM System One located at the Cleveland Clinic to run the experiments. Due to the inherent noise in current quantum devices, we anticipated challenges in model convergence and performance stability. We took a warm start approach together with error mitigation techniques including Dynamic Decoupling and Pauli Twirling along with increased number of shots. We found there to be a linear increase in model runtime as a function of number of shots for both our 20 qubit as well as our full peptide 46 qubit model, with experiments using 20k shots completing within reasonable runtimes of approxiamately 15-17.5s (Fig. 2E). For this warm start approach, the initial weights to be used in the hardware experiments were extracted from the noiseless statevector simulation experiments, and the learning continued on the QPU, as depicted in Fig. 2D. By analyzing the loss values produced after each forward pass of the circuit, we roughly determined when the model has converged to some optimal set of weights. We observe that the model converges after approximately 660 iterations, indicating a local minimum of the loss function that best satisfies our MHC-binding prediction model. To test proper behavior on the hardware, we then warm start the same model, using the weights previously calculated at the final iteration in the simulation, with the quantum hardware as the backend for an additional 200 iterations. We found that the resumed hardware results aligned with the simulated behavior observed in this test case, achieving a F1 score of approximately 0.80. This workflow based on testing on hardware after training on the simulator allowed us to further assess the error mitigation strategy and confirmed the hardware was operating as expected.

### Benchmarking QCNN binding prediction

To evaluate the performance of our QCNN model on predicting MHC-binding, we benchmarked it against two different classical Convolutional Neural Networks (CNNs) using identical training and testing splits as well as identical binary encoding methods (Table 2). The first CNN baseline was designed to reflect the current limitations of the quantum devices. Specifically, it was restricted to operate under similar architectural constraints that limit the number of trainable parameters. On the quantum side, our QCNN had parameter counts that range from 21 in low input settings (i.e. 4 qubit experiment) to 399 in the full peptide-coded input (i.e. 46 qubit experiment). In contrast, typical classical CNNs, when applied to the full peptide encoding, often involve thousands of trainable parameters, reflecting significantly greater model complexity. Therefore, to ensure a fair comparison, we built a dynamic classical CNN that similarly matched the number of trainable parameters to the QCNN. Additionally, we constructed a model that reflects current ML standards; therefore, our second CNN baseline was unrestricted and contained thousands of trainable parameters.

**Table 2.** Difference in learnable parameters between the QCNN with the classical CNN models.

| Positions | QCNN | CCNN Ltd. | CCNN Large |
| --- | --- | --- | --- |
| BA RANK | 21 | 25 | 55873 |
| 3 Positions + BA RANK | 54 | 125 | 55873 |
| 4 Positions (QRAC 2:1) | 81 | 125 | 55873 |
| 4 Positions (Basis) | 168 | 125 | 55873 |
| Full Sequence | 399 | 329 | 55873 |

**Table 3.**
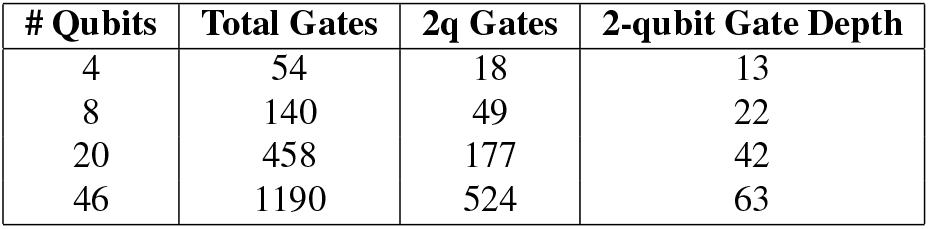
QCNN circuit complexity across different qubit counts. The table reports the total number of gates, the number of 2-qubit gates, and the 2-qubit gate depth (i.e. the number of layers containing at least one 2-qubit gate) for various input sizes.

After developing both of the classical CNN baselines, we trained and tested each of the models on identical train and test datasets as the QCNN model in addition to using the same binary encodings of the input. As the BA RANK determines how likely a peptide will bind to an MHC complex, we tested each model with an input of strictly BA RANK without any amino acids to give a baseline for the effectiveness of each model. Additionally we tested both models using an input of peptide positions 2, 5, and 9 with BA RANK, as well as peptide positions 2, 4, 5, and 9 without BA RANK. When comparing results across the three binding prediction experiments (Table 1), it is evident that the classical CNN constrained to match the QCNN’s parameter count performs worse, especially in capturing complex patterns within the data (e.g. positions 2, 4, 5, and 9 without BA RANK). We then compared the prediction outputs of the QCNN and classical models against NetMHCpan’s BA RANK labels as a baseline (Fig. 3). By analyzing the predicted labels across all four approaches, we observe that although the overall performance appears to be similar, the models still assign different predictions to individual samples. This suggests that the QCNN model is learning patterns or decision boundaries within the data that are different from those learned by the classical models.

**Figure 3.**
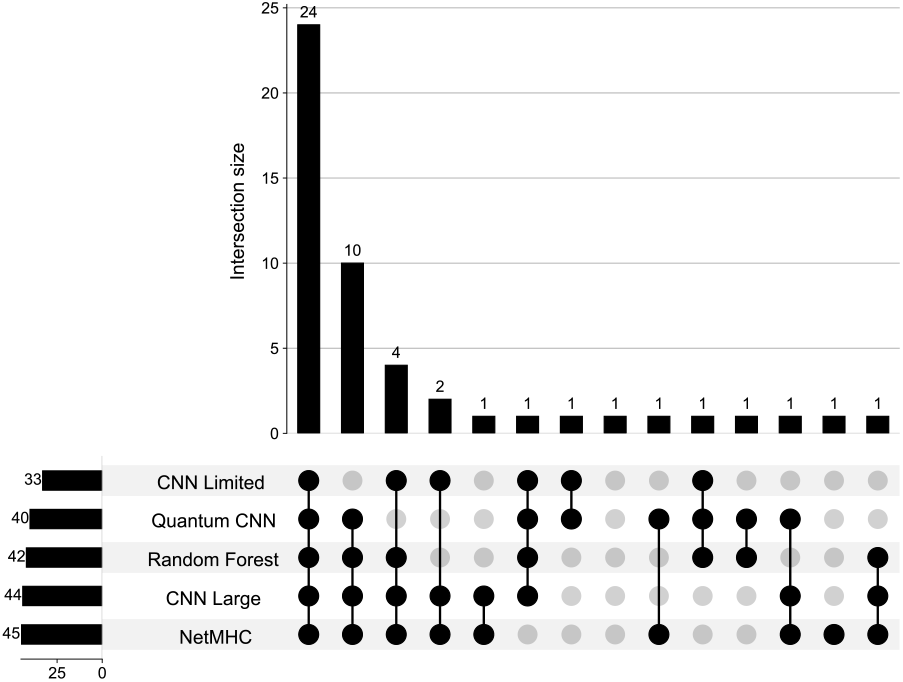
Benchmarking QCNN against classical methods for MHC binding classification. Upset plot depicting the differences in predictions between the QCNN, two classical CNNs, and random forest model consisting of for testing dataset consisting of 50 peptides. Training for the models included strictly peptide positions 2, 4, 5, and 9. Additionally included in NetMHCpan’s BA RANK predictions for binding. Light gray nodes indicate incorrectly labeled samples for the model while black nodes indicates correct MHC binding predictions. The intersection plot above shows the number of samples shared between each of the models.

### Immunogenicity prediction accuracy can be over inflated when including non-binders

With successful modeling of MHC binding via QCNN we next considered immunogenicity predictions. Immunogenic peptides must be both bound by the MHC and recognized by a T-Cell receptor (Fig. 4A), making it challenging to capture both experimentally and computationally. In our recent work^9^, we developed prior screens to capture both binding and T cell receptor recognition, and we observed that the input of the data could drastically alter the accuracy of the model used for immunogenicity prediction (Fig. 4B). When using NetMHC to predict immunogenicity, the predictive performance (AUC) is 0.79 when including those peptides that do not bind MHC and those that do bind MHC (Fig. 4B). However, if only those peptides experimentally confirmed to bind to MHC are used, then the performance of the model drops to 0.55 (Fig. 4B). This is a result of the class imbalance in the datasets, which are skewed with few numbers of immunogenic peptides compared to the number of the non-immunogenic peptides in the full dataset of non-binders and binders. Thus, a the model can achieve a high accuracy when classifying most peptides as negative. Given this observation, when we approach modeling immunogenicity with quantum methods, we primarily focus on training with MHC binders to capture features of immunogenicity.

**Figure 4.**
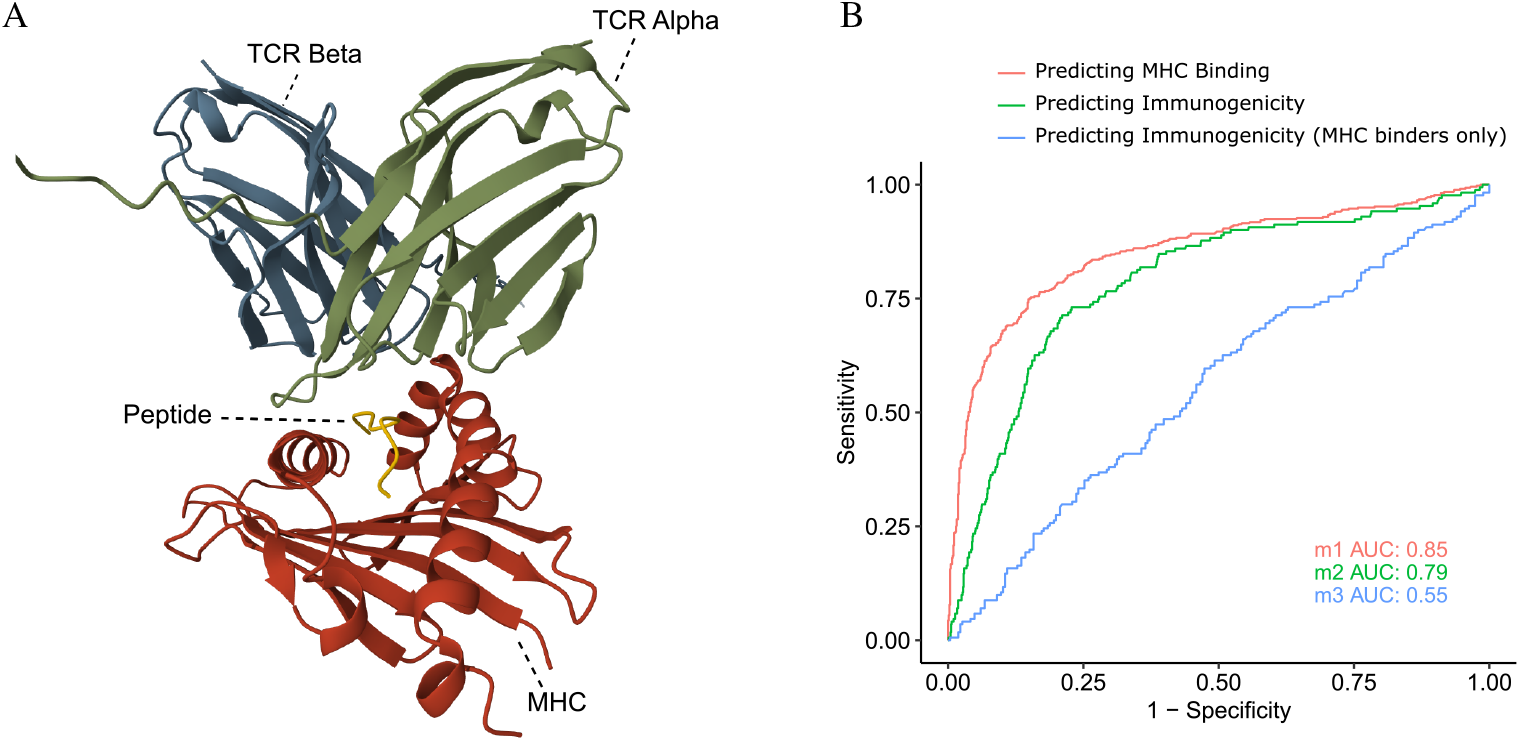
Benchmarking NetMHCpan for MHC binding and immunogenicity prediction. A) NetMHCpan benchmarking using Alban et al., 2024 Nature Medicine dataset^9^. which includes 1,453 neoantigens, 196 of which are experimentally confirmed as immunogenic. NetMHCpan demonstrates effective MHC binding predictions as it achieves a high AUC of 0.85. It additionally appears to accurately predict immunogenicity when including those peptides that are known MHC binders and non-binders (AUC score of 0.79). However when restricted to the appropriate context of only confirmed MHC binders, the AUC drops to 0.55, highlighting a key challenge in the field: predictive performance is highly dependent on dataset composition. B) Immunogenicity model depicting the ability of a peptide presented on MHC to be recognized by a T cell receptor (TCR), triggering an immune response.

### QCNN prediction of immunogenicity

Expanding our quantum efforts to immunogenicity prediction we first encoded each peptide position encoded into a five-bit binary vector representing the amino acid at that position. We once again utilize the noiseless statevector so that the initial results reflect the circuit’s intrinsic behavior, free from hardware noise, letting us verify that the model functions as intended.

In order to identify the positions most predictive of immunogenicity, we conducted a feature parameter search by evaluating which combinations of four positions produced the highest model performance. While simulators are effective at modeling a low number of qubits, scaling to larger experiments, particularly those greater than 20 qubits, results in exponentially increased runtimes. In order to reduce the scale of each of the experiments, classical encodings were therefore mapped to quantum states using QRAC 2:1, resulting in a manageable 10 qubit circuit.

To ensure our model was properly trained, we increased our dataset to include 300 training and 100 testing samples, maintaining balanced class distributions across both. Initially, we ensured the split contained an equal proportion of confirmed immunogenic peptides; those that were not labeled as immunogenic were able to be both MHC binders and non-binders. After running a feature parameter search, we found the model with encoded peptide positions 1, 3, 4 and 9 was most effective in predicting immunogenicity, achieving a F1 score of 0.77. Further analysis revealed that position 9 consistently appears in top-performing models, highlighting its potential importance (Ext. Data Fig. 3B). When averaging performance across all tested positional combinations, we observed a lower F1 score of 0.64, which is expected given that only select positions meaningfully contribute to immunogenic response. We additionally evaluated the same sets of residue positions with the inclusion of BA RANK, as this feature is expected to enhance the model’s ability to filter out non-binding peptides. Using the same training and testing splits as in the previous experiments, we observed a decrease in the F1 score (0.73) when including BA RANK with positions 1, 3, 4, and 9. When analyzing overall performance across all position sets, the inclusion of BA RANK generally does not help the model as it achieves a similar average F1 score of 0.64 across all models.

In order to ensure the model was not overfitting to patterns specific to MHC binding predictions, we developed additional data splits using samples that are confirmed to bind to a MHC complex. We subsample this data subset to include 300 training samples and 100 testing samples. Next we use a similar approach as before to identify those positions that are most effective in predicting immunogenicity by iterating through different splits of four positions. Among all tested combinations, the model using positions 3, 5, 6, and 7 achieved the highest F1 score of 0.67. Further analysis of the top-performing models revealed that positions 5, 6, and 7 consistently appeared to be key features (Ext. Data Fig. 3C), which is consistent with positions known to interact with the T cell receptor. When averaging the performance across all tested positional combinations, we observed a lower F1 score of 0.51. As previously mentioned, this reduction is expected since only select positions are helpful in determining immunogenic response. Also as expected, including BA RANK in the input sequence yielded minimal improvement in model performance. The best-performing model once again included positions 3, 5, 6, and 7, achieving a F1 score of 0.70. When averaging performance across all positional combinations, we observed a slight increase, with an average F1 score of 0.52. This marginal gain was to be expected, given that all peptides in the training and test sets are already assumed to be binders.

### Immunogenicity prediction with full length peptide model

As with our binding prediction experiments on QPUs, we are now able to use a basis feature map, thereby avoiding potential information loss from the QRAC 2:1 feature map, as we are no longer constrained to low numbers of qubits by the quantum simulator. First we begin our quantum hardware experiments by examining positions 4, 6, 7, and 8 with BA RANK included in the input. For data input we chose a smaller subset of 150 training samples and 50 testing samples that include both binders and non-binders as we were limited by both runtime and system restrictions. After execution, we observe that the model achieves a F1 score of 0.74. Following these results we also tested the model including positions 1, 3, 5, and 9 as well as BA RANK and observed a similar F1 score of 0.75. With confidence from both simulator and initial hardware experiments, we decided to encode the full peptide as input for our model. Using binary encoding with a basis feature map, our input translates to a 46 qubit experiment (five qubits for each amino acid position in addition to an additional padding qubit needed for the QCNN pyramid circuit architecture). As this is a much larger experiment than previous efforts, we needed to ensure proper error mitigation techniques to prevent the noisy environment from interfering with the results.

For initial testing, we substituted the extra padding bit with a bit indicating BA RANK, keeping the circuit size the same. Previous efforts confirmed that BA RANK significantly decreased the number of iterations needed for the model to converge (Fig. 2A). Therefore to test error mitigation techniques for the 46 qubit experiment, we did an initial BA RANK test. We found the runtime of the circuit was comparable between our prior 20 qubit hardware experiment and this 46 qubit, up to 20k shots (Fig. 2E), therefore we selected 20k shots for further 46 qubit experimentation given that our smaller 20 qubit circuit was able to converge at this setting (Fig. 2D). We then ran our full peptide experiment without BA RANK. After approximately 350 iterations, the model was able to converge and achieved an F1 score of 0.70 (Fig. 5). After iteration 400, the loss began to rise indicating instability that may be attributable to increased noise sensitivity of this 46 qubit circuit model because of changes in device noise profile over the duration of training lasting multiple days.

**Figure 5.**
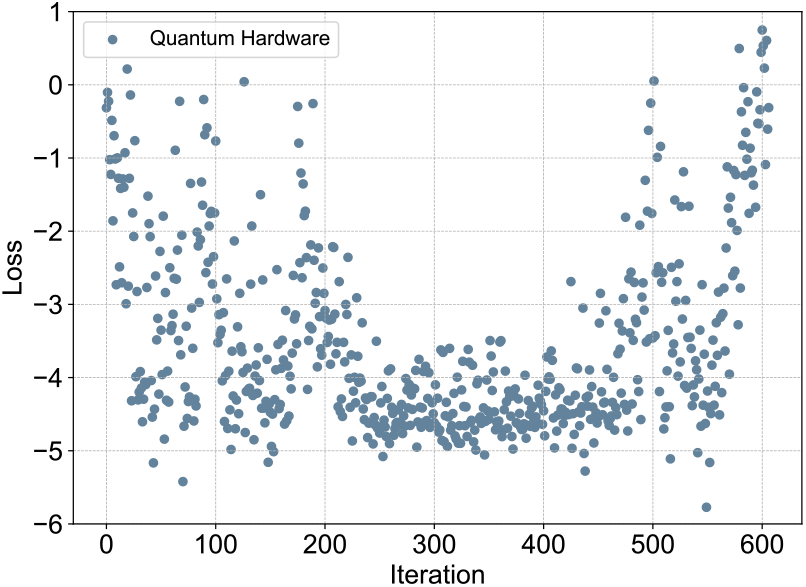
Loss curve for immunogenicity prediction using full peptide sequences. The model was trained and tested on 150 and 50 peptides, respectively, with balanced class distributions. To mitigate quantum noise, the experiment used 20,000 shots per circuit along with Pauli Twirling and Dynamic Decoupling. Despite these techniques, the plot reveals a period of instability, particularly between iterations 400 and 600, likely due to increased noise sensitivity of the larger 46 qubit circuit.

### Benchmarking QCNN immunogenicity prediction performance

Similar to our binding prediction models, we were interested in benchmarking our current quantum results with classical ML models, such as CNNs and random forests. Here we selected random forest as a secondary benchmark as random forest is one of the simplest machine learning architectures and has been a standard benchmark in drug discovery applications for over two decades. We first created a dynamic CNN to model the lower number of parameters within our experiments (21 parameters for a small experiment to 400 parameters for the full peptide encoded sequence). Additionally, we created a model that allowed for more parameters to capture more complex patterns. Finally we developed a random forest model. For each of the models we trained and tested on the larger 300 train, 100 test datasets and included data subsampled from those peptides that were confirmed to be binders only as well as a mix of both binders and binders. Additionally, we tested each with and without the presence of BA RANK to determine the importance of the added feature. The results are summarized in Table 4 and 5.

**Table 4.** Summary of immunogenicity predictions by classical benchmarks against the QCNN model for the datasets containing both MHC-binders and non-binders. Classical models include the limited parameter classical CNN (CNN (ltd.)) to model the low number of parameters in a QCNN, the unrestricted parameter classical CNN model (CNN (large)), and a random forest model (RF). All results are measured as an F1-Score. The full peptide model was run on a PFTQD using a smaller 150 training split due to hardware restrictions (*).

| Positions | QCNN | CNN (ltd.) | CNN (large) | RF |
| --- | --- | --- | --- | --- |
| BA | **0.73** | **0.73** | **0.73** | 0.68 |
| 1,4,5,9 | **0.71** | 0.33 | 0.67 | 0.64 |
| 1,4,5,9,BA | 0.71 | 0.50 | **0.72** | **0.72** |
| 4,6,7,8 | 0.49 | 0.39 | 0.33 | **0.52** |
| 4,6,7,8,BA | 0.54 | 0.35 | 0.68 | **0.72** |
| 4,6,8,9 | 0.70 | 0.35 | **0.71** | 0.68 |
| 4,6,8,9,BA | 0.74 | 0.55 | 0.71 | **0.77** |
| Full peptide | **0.70** * | 0.60 | 0.65 | 0.66 |

**Table 5.** Summary of immunogenicity predictions by classical benchmarks against the QCNN model for the datasets subset to only MHC-binders. Classical models include the limited parameter classical CNN (CNN (ltd.)) to model the low number of parameters in a QCNN, the unrestricted parameter classical CNN model (CNN (large)), and a random forest model (RF). All results are measured as an F1-Score.

| Positions | QCNN | CNN (ltd.) | CNN (large) | RF |
| --- | --- | --- | --- | --- |
| BA | 0.54 | 0.33 | 0.54 | **0.62** |
| 3,5,6,7 | 0.67 | 0.42 | **0.68** | 0.65 |
| 3,5,6,7,BA | **0.70** | 0.35 | 0.65 | 0.69 |

### Q-CHIPP predicting response to immunotherapy

To assess the biological usefulness of our prediction model, we next applied it to a cohort of peptides derived from patients with lung cancer treated with immunotherapy, to determine if it could be used to separate response to therapy^29^. This technique has previously been used to evaluate immunogenicity models^7^. Here we select only those patients with HLA-A02:01 (n = 111) and use a 9 amino acid long sliding window around each somatic mutation to generate 209,889 peptides. The peptides within the data set were not experimentally screened for immunogenicity; instead predictions from our model were used to stratify the patients based on their predicted burden of immunogenic antigen peptides. For this we effort, we developed a combination of our best performing models, which we named Quantum HLA Immunogenicity Peptide Prediction (Q-CHIPP). Q-CHIPP is made up of two models: one trained on the IEDB data set to predict binding using peptide positions 2, 4, 6 and 9, and the second model is designed to predict immunogenicity with input of peptide positions 3, 5, 6, and 7. Importantly, the immunogenicity model is trained exclusively on peptides confirmed to bind to the MHC, ensuring that the features learned are specific to immunogenicity rather than confounded by MHC-binding. This dual-model framework allows Q-CHIPP to integrate distinct but complementary signals, binding potential and immune recognition, to improve overall immunogenicity prediction performance. To aggregate the two models for Q-CHIPP, we predict a given peptide to be immunogenic if both the binding and immunogenicity models independently classify it as positive. Using both the individual models as well as our Q-CHIPP model, all 209,889 peptides were evaluated and the total number of predicted immunogenic peptides were aggregated for each patient to estimate the immunogenic neoantigen load.

To evaluate the predictive utility of our model, we bench-marked its performance against NetMHC’s binding affinity rank. As mentioned previously, BA RANK represents a ranking score of the likelihood that a peptide is bound to an MHC molecule, with lower values indicating a stronger binding potential. A conventional threshold of 2.0 was used to classify each peptide as a potential binder and these total predicted counts were aggregated for each patient, similarly stratifying the patients based of their total predicted counts. We additionally benchmarked our Q-CHIPP model with two classical CNN’s as described before; the first limiting the number of parameters in the model to closely match the QCNN model (CHIPP-limited), and the second being unrestricted in the number of parameters (CHIPP-large). Both classical CNNs used identical training splits and shared the same combination of individual models: one pre-trained for MHC binding using positions 2, 4, 5, and 9, and the other for immunogenicity prediction using positions 3, 5, 6, and 7.

On studying the results of each of the survival analyses, we found that while all models were significant in their ability to differentiate between survival, Q-CHIPP had the strongest p-value (p=0.0085) (Fig. 6A-C). Additionally, all models out performed NetMHC alone (p=0.0275) (Fig. 6D). Comparing our Q-CHIPP and CNN methods to more refined and unconstrained methods like Random Forest demonstrated similar performance for both binding and immunogenicity models, but as expected the Random Forest model is still able to slightly out perform Q-CHIPP (p=0.0052) (Ext. Data Fig. 4A-I). Importantly, in the most fair benchmark of these methods Q-CHIPP outperformed the individual QCNN models for Binding and Immunogenicity (Fig. 6A-C). This indicates that the comminatory approach between two distinct models may be more sensitive to identifying immunogenic antigen load. Additionally, analyzing the differences in predictions we see that Q-CHIPP identifies 25,542 peptides that are not picked up by any of the other bench-marked methods including NetMHC, random forest, classical CHIPP-large, and classical CHIPP limited (Fig. 6E). To better understand what features Q-CHIPP is selective for we next looked at the predicted positive/negative amino acid ratios per position for Q-CHIPP, CHIPP-large, and CHIPP limited (Fig. 6F-H). These ratios demonstrated that the Q-CHIPP model is easily picking up strong Amino acid signals at both binding positions 2,9 as well as immunogenicity positions like 4,and 7 (Fig. 6F). Particularly, position 7 has strong enrichment of Alanine, Aspartic Acid, Lysine, Methionine, Threonine, and Tryptophan. These features are not as selected for at position 7 in the CHIPP limited model or the CHIPP large model. In contrast the CHIPP limited and CHIPP large model both had a strong bias for peptides with Leucine at position 2, which may indicate the model is biased toward features of binding and not immunogenicity (Fig. 6F-H).

**Figure 6.**
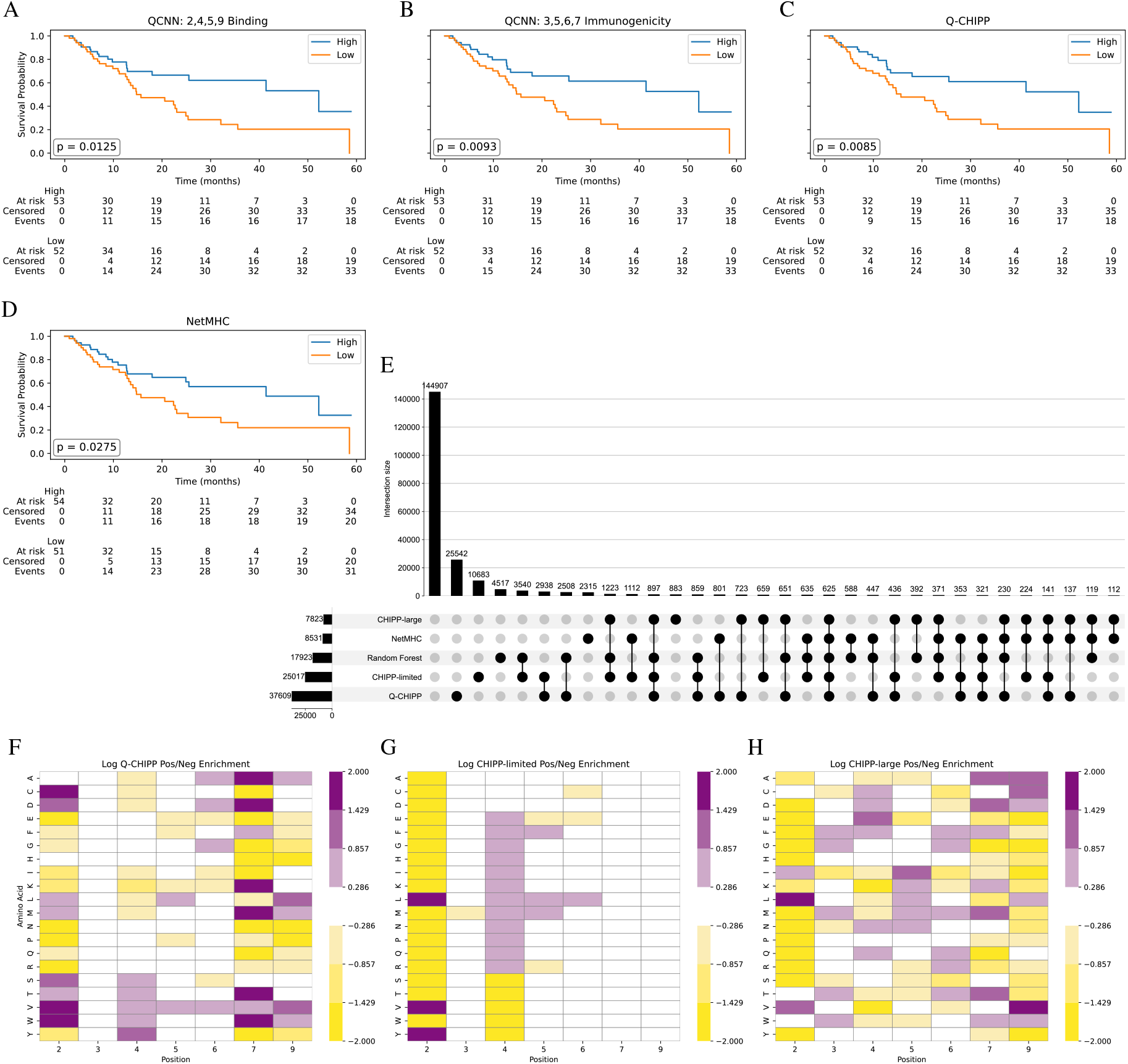
Patient survival predictions for A) QCNN binding predictions from pretrained model on IEDB dataset using positions 2, 4, 5, and 9, B) QCNN immunogenic predictions from pretrained model on IEDB MHC-binders only dataset using positions 3, 5, 6, and 7, C) Q-CHIPP model composed of the aggregation of models A and B, and D) NetMHC values using threshold of 2.0 to classify between likely binders and non-binders. E) Differences in positive labeled predictions between the Q-CHIPP model, benchmark models CHIPP-limited, CHIPP-large, and random forest, in addition to NetMHC. Light gray nodes indicate negative labeled samples for the model while black nodes indicates positive predictions. The intersection plot above shows the number of samples shared between each of the models. F) Q-CHIPP, G) CHIP-limited, H) CHIP-large. Positional amino acid enrichment calculated as the ratio of amino acid frequencies at each position in positively predicted samples versus negatively predicted samples.

## Discussion

Immunologic recognition and removal of non-self antigens is based on the protein-protein interaction of MHC, antigen, and T cell receptors, and while more complex systems have been computationally modeled, this problem remains challenging. The heterogeneity of HLA alleles is estimated to be greater than 26,000 world wide and the possible combination of unique T cell receptors is estimated to be 10^20^ or more, creating a significantly more complex problem than simply modeling the interaction of 3 proteins^30^. In addition to the large heterogeneity among T cell receptors and MHC, cancer neoantigens only differ from self-peptides by one amino acid, demonstrating the incredible evolution of immune specificity that can be generated by random combinations of T cell receptors. These challenges along with the patient specificity of each interaction makes them not only difficult to predict but also challenging to screen for. Due to limited predictive tools and the high cost of experimental screens, the number of available neoantigens for model training remains small. To address this, many groups have turned to meta-analyses that aggregate data from multiple sources. We have also adopted this strategy by combining our own recently developed neoantigen screening data from a lung cancer clinical trial of immune checkpoint therapy. In our screen, we tested both MHC binding and T cell recognition separately, which is unique as many studies perform this in a single step to reduce costs at the expense of limiting the information we can learn about determinants of binding vs T cell recognition. We begin our paper demonstrating the importance of this information and how the current standard in the field, NetMHC, appears to predict immunogenicity when both MHC binders and non-binders are present in the test data, however once non-binders are removed, NetMHC is no better than chance. With dataset training size representing a primary hurdle we sought out the use of QML as an alternative to classical ML to model immunogenicity.

In choosing a quantum technique, QCNN, similar to current neoantigen modeling techniques, we had a similar conceptual framework for encoding each amino acid and position into qubits. Previously we demonstrated that these sequence based approaches lose the dynamic information that can be captured with molecular dynamic simulations, but the current limitations of quantum hardware do not allow us to conduct such large scale dynamic and structural based models. As quantum hardware scales it is our hope that someday we may be able to use quantum chemistry approaches to model the full HLA binding pocket, peptide, and TCR, but it is currently estimated that we would need approximately 600 qubits to model just the key atoms involved in the interaction. Instead, we focused first on the near term quantum solution of using a QCNN model to predict both MHC binding and immunogenicity and found that the QCNN is able to predict MHC binding with high accuracy using a small training dataset. Using only four amino acids of the total peptide, we were able to scale our model onto quantum hardware and run various models scanning peptide positions, with QCNN showing the strongest prediction accuracy with anchor residues 1,9 and the TCR contact residues 4,5,6. While this biology is well understood and described, it is interesting that the QCNN approach can identify these with a relatively small training dataset. Initially when comparing directly to CNN, we observed no major differences in performance, but when CNN is limited to the same size parameter space as QCNN, due to QCNN hardware restrictions, we see that QCNN has a clear advantage over CNN. Additionaly, we noted that the QCNN and CNN models are selecting for different amino acids at each position, demonstrating they are learning differently. We feel that this provides evidence that as the QCNN is able to learn with an order of magnitude fewer parameters and suggests that as quantum hardware scales to support more complex, expressive circuits, it may greatly outperform classical methods at this task.

After developing our model to predict binding, we next focused on predicting whether the neoantigen triggered an immune response. Initial testing reveals that when MHC binders and non-binders are present in the training, the model appears to be performing relatively well, achieving an F1 score of 0.71 for top performing models. However, when analyzing which peptide positions correspond to the best predictive models, we see that position 9 has a significant presence. Comparing this to the MHC binding predictions, we see a similar trend where peptide position 9 appears to be important in achieving more accurate results. This indicated to us that while the model appears to be performing well on the test set, in reality the model is likely capturing simpler patterns of predicting those peptides that bind to the MHC. To account for this, we tested the model using a subset of only peptides that are MHC binders. We saw that while the model performance decreases, peptide positions 5 and 6 become key features in top performing models, aligning it with the biological expectations.

When analyzing both binding and immunogenicity predictions, we found that while classical benchmarks (NetMHC, classical CNN, random forest) may outperform our quantum classification model when unrestricted to equal parameter size, the QCNN was able to capture significant trends within the data given a much lower number of samples, subsample of features (four features for initial models), as well as a limited number of trainable parameters used within the model, indicating the potential efficiency and power behind QML. When comparing the quantum model with the classical CNN, we found that when the two models had similar number of parameters and identical training and testing datasets, the quantum model was able to capture much more complex trends within the data than the classical counterpart. We further expand on these findings by developing a combinatorial model using two models: one trained on binding and one trained on immunogenicity within a know set of MHC binders. Combinatorial approaches were suggested by recent consortium studies^7^ and we also explored this concept in our recent work^9^, but this represents the first large-scale effort to use this approach with Quantum computing and to our knowledge classical methods also. With this approach we showed that the predicted neoantigen load from any combinatorial model we developed out performed the single binding or immunogenicity trained model when predicting response to immunotherapy. Future efforts should explore the use of these combinatorial models by screening top candidates.

While there are many exciting points throughout our work it is important to recognize that quantum hardware today is inherently noisy, which can limit the model’s performance and increase runtime due to the added complexity of error mitigation strategies. However, as quantum technology advances, particularly in error correction, qubit coherence, and gate fidelity, we expect the performance of quantum models like the QCNN to become both more accurate and computationally efficient. We saw within the given experiments that for low number of qubits (i.e. less than 20), the noiseless statevector simulators are more computationally attractive without the need for the error mitigation techniques. However, with each additional peptide encoded into the circuit, the runtime exponentially increases on the noiseless simulators. Therefore after 4 encoded amino acid positions, we are forced to transfer the model to the PFTQD. With these caveats we were still able to run a large scale 46 qubit experiment incorporating full length peptides trained to predict immunogenicity. As we described the model initially converged and yielded an F1 of 0.71, but then later picked up noise again as iterations continued. We noted that while this happened several of the qubits in our circuit picked up a high error rate, and may have contributed to the loss of convergence. In addition to these technical aspects, our models currently predict binary outcomes only, and do not provide a ranking method like other tools in the field. As a future technique we plan to develop similar approaches with a continuous rank via regression instead of binary outputs so that we can better rank peptides for screening and evaluation against other tools in the field.

Our results suggest that the quantum convolutional neural network already offers meaningful advantages for neoantigen modeling even on today’s limited hardware. We highlight that the model is able to learn the biologically relevant anchor and TCR-contact residues with a relatively small training size and retain predictive power when classical models are under the same parameter budget. Although current noise levels, qubit limits, and circuit restrictions confine our experiments to limit the training size, number of encoded positions, and number of learnable parameters, there is clear separation between the QCNN and classical counterparts, suggesting the QCNN is able to learn different patterns within the data. Looking ahead, the continued evolution of quantum hardware, including increased qubit systems, improved coherence times, and advanced error correction, will enable more expressive and scalable quantum circuits, bringing us closer to modeling the full complexity of neoantigen modeling. Ultimately, quantum machine learning holds significant promise not only for advancing immunogenicity prediction, but for also contributing to the broader goal of precision immunotherapy, tailoring vaccines and T cell therapies based on individual patient profiles.

## Methods

### Data Sets

Immunogenicity datasets and were utilized form multiple sources including IEDB^31^, Tesla consortium data^7^, PRIME model training data^8^, and our own internal data set from lung cancer neoantigen screening^9^. IEDB was primarily utilized due to the large number of immunogenic antigens with the HLA-A02:01 allele, while the binding vs non-binding comparisons were focused on our internal dataset from^9^ due to the experimental design that allowed for binder vs non-binder analyses. Additionally, we used whole exome sequencing data from^29^ to predict neoantigens from for survival analyses. All raw dataset IDs are identified in data availability section.

### Data Encoding

After curating the respective datasets, each cohort was encoded into the appropriate input format for the model. Each of the 20 amino acids are mapped to a unique integer that is then used for either one-hot encoding or binary encoding. This helps to avoid introducing false ordinal relationships through numerical encoding, over the nine amino acid positions of the neoantigen peptide sequence. With binary encoding (`category_encoders.BinaryEncoder()`, we convert each integer into a distinct five-bit binary vector. As 2^5^ = 32 possible combinations, this is sufficient to represent all 20 amino acids. Additionally, it is worth noting that each of the 9-positions were zero-based indexed when encoded. Before instantiating each QCNN model, an additional layer of data encoding is implemented to ensure proper input for quantum feature maps. Quantum circuits return expectation values in range [-1, 1], so the ground-truth labels were remapped from conventional {0, 1} encoding to {−1, 1}. Additionally, each of train and test datasets were subset to only include only those features specified for the given model (ex. `[motif1, motif4, motif7, motif8]` where motif represented the zero-indexed peptide position).

For each peptide, NetMHCpan’s binding affinity (BA) rank was computed, and this value was converted to a binary value using a threshold of 2.0 to classify likely MHC binders (assigned 1 for ranks *<* 2.0) and non-binders (assigned 0 for ranks ≥2.0). Additionally, each of our truth labels for MHC binding and immunogenicity, confirmed via experimental screening, were encoded similarly as binary values indicating confirmed binding or immune response. The QRAC 2:1 and 3:1 feature maps require a multiple of two or three classical bits, respectively, for every quantum bit. Additionally, the QCNN pyramid circuit design requires an even number of qubits after being mapped. Therefore to account for these we added padding columns which were simply zeros encoded, allowing us to meet the respective requirements for an experiment.

### Dataset Splits

To ensure reproducibility, all random operations during dataset preparations were performed using a fixed random seed (1). For each classification task, either MHC binder versus non-binder or immunogenic versus non-immunogenic classification, the dataset was first divided based on their corresponding binary label into the two class groups. From each class, an equal number of samples were randomly selected using `pandas.sample` with `random_state=1`. Each of our datasets consisted of either 150 or 300 training samples depending of the experimental setup. These sizes were selected to balance model performance with the limitations of circuit executions and runtime on quantum hardware. The selected subsets were then concatenated and passed to `sklearn.model_selection.train_test_split` with a 75:25 train-to-test ratio. Stratification was applied based on the label to preserve class distribution in both sets, and the same random seed (1) was used to maintain consistency. This approach was used to ensure reproducible and balanced splits that were appropriate for the quantum devices.

### QCNN Model Architecture

To construct the QCNN we use Qiskit (v1.4.2), an open-source framework for developing quantum computing applications. One of its key strengths lies in its focus on hardware-agnostic development, enabling the possibility to run on different quantum hardware platforms. Our QCNN circuit was developed using the available QCNN circuit as part of the open source Qiskit Machine Learning (v0.7.2) as the basis for our implementation with significant modification to increase circuit efficiency by minimizing the number of SWAP gates needed through a pyramid layout of the SWAP gates.

We begin our QCNN circuit with a feature map responsible for translating the encoded classical input to qubits within the quantum space. Several different options are available for the feature mapping. Z and ZZ feature maps utilize the `ZFeatureMap` and `ZZFeatureMap` from `qiskit.circuit.library` package. Here we apply a Z-axis rotation (RZ gates) and entangling operations. Quantum random access code (QRAC) mappings allow for the compression of classical information where 2 or 3 classical bits are encoded to a single qubit. Additionally available is the basis feature map which applies single-qubit Pauli rotations (RX) to encode classical data. Here we primary utilize the QRAC 2:1 for smaller scale experiments on the simulators as well as the basis feature mapping.

We add to this circuit our QCNN pyramid ansantz. The ansatz is composed of multiple convolutional and pooling layers placed one after the other. Starting with an initial *n* qubits after the feature map ansatz, the convolution layer is implemented as a sequence of two-qubit unitaries called the convolution unitary on alternating nearest neighboring pairs of qubits as shown in Ext. Data Fig. 2. Each convolutional unitary is composed of 3 CNOT gates and 3 parametric single-qubit gates. This is followed by the pooling layer formed with 2-qubit pooling unitary. Each pooling unitary is composed of 2 CNOT gates and 3 parametric gates acting on pairs of qubits (*q*_*i*_, *q*_*i*_ + 1) where *i* starts with qubit 1 or 2 based on whether total number of active qubits are a multiple of 4 or not for that layer. At the end of each pooling layer, we discard qubits symmetrically from the top and bottom of the register, retaining the central 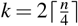 qubits. A series of SWAP gates are additionally placed between qubits to align the central qubits into neighboring positions. We repeat this process with next layer composed of 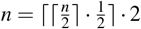 qubits. This is repeated until we have *n* = 1 qubit remaining. This qubit is then measured in the Z basis.

To execute our quantum circuits on quantum hardware we use `QiskitRuntimeService` (v0.30.0) with the estimator primitive. Simulated experiments are performed using `AerSimulator` (v0.15.0) backend from Qiskit to model ideal, noiseless behavior on smaller scale experiments. For hardware-based runs, we employ `ibm_cleveland`, an Eagle R3 processor featuring 127 qubits. When utilizing a PFTQD such as `ibm_cleveland` it is crucial to ensure the noisy devices give accurate results. We therefore employ several error mitigation and suppression techniques including Dynamical Decoupling and Pauli Twirling via the estimator primitive. We additionally adjust the number of shots, *s*, which reruns each circuit *s* times to ensure accuracy. For smaller experiments (20 qubits) we found *s* = 5, 000 was sufficient to get accurate results, however for our larger (46 qubit) experiments, we utilized *s* = 20, 000. When submitting to a QPU, we specify an initial layout of the qubits to optimize qubit quality for our ansatz. We represent the QPU’s qubit connectivity as a graph with edge weights based on the connection error between qubits and node weights the qubit readout error. Nodes and edges with error levels above a user-defined from threshold (readout_error_threshold=0.12, edge_error_threshold=0.2) are removed from the graph. All paths between of specified length, a function of the required number of qubits, are identified and the median readout error of the entire path as well as the central four qubits are calculated. These central four qubits are important as they are the last remaining qubits of the circuit and therefore require the longest coherence time, particularly the final qubit. The optimal path is selected by first filtering for the top 5% of paths by their median overall readout error, and then selecting from this subset the path with minimal median readout error for the central four qubits. This path is used for the initial layout of the circuit. It is important to note that if the model is resumed later using previously calculated weights to specify this initial layout for consistency in circuit mapping and model behavior. After specifying the initial layout of the qubits, we transpile the circuit to ensure capability with the backend. We utilize a pass manager from Qiskit’s `generate_preset_pass_manager` with optimization level of 1, as higher optimization levels are not required given our earlier search for the optimal initial layout.

### Training

For each model, a set seed was initialized for packages `qiskit_algorithms` and `numpy` (49) to ensure reproducibility of the experiment. To build our parameterized quantum circuit integrated with our estimator, we utilize the `EstimatorQNN` from `qiskit_machine_learning` (v0.7.2). We utilize Simultaneous Perturbation Stochastic Approximation (SPSA) optimization (`qiskit_algorithms.optimizers`, v0.3.0), which is a gradient-free optimization algorithm useful when in noisy environments. The optimizer takes a learning rate and perturbation functions as parameters. For the learning rate we use `spsa.powerseries` with `eta=learning_rate_a` (user-specified parameter), `power=0.602`, and `offset=0`.

Similarly for the perturbation function we use the `spsa.powerseries` with `eta=0.2`, `power=perturbation_gamma` (user-specified parameter), and `offset=0`. If the `learning_rate_a` and `perturbation_gamma` parameters are not provided, an initial set of iterations will occur to calculate the ideal parameters. However with extensive runtimes on the quantum hardware devices, we found it was best to preset these parameters to reduce runtime. We found using `learning_rate_a=0.06` and `perturbation_gamma=0.101` was ideal for our model.

In order to handle training and testing of our neural network, we utilized the `NeuralNetworkClassifier` from `qiksit_machine_learning` (v0.7.2). Here we give it the developed `EstimatorQNN`, `cross_entropy` loss function from Qiskit, and the SPSA optimizer. We additionally provide it any previously calculated set of weights if we choose to warm-start the model and supply it with a callback function used to keep track of the current weights and loss values.

### Model evaluation

After training, we evaluate each model on an unseen dataset according to several metrics. We computed confusion matrices for each model, analyzing the number of true/false positives/negatives. As our we were using balanced train and test splits for binary classification, we utilized the F1 score to analyze the overall performance at predicting the MHC binding or immunogenicity. Additional metrics included accuracy, ROC AUC, precision, and recall; however most of these metrics reported very similar scores due to the even class distribution used for training and testing. Additionally, to measure the models performance during training, we analyzed the objective function values at each iteration training. As mentioned, we utilize the callback graph within our `NeuralNetworkClassifier` which saves the weights and objective function values at each iteration of training. We can therefore plot these loss values and analyze if the model appears to be converging, indicated by a steep downward trend and then flattening to a steady constant value. These metrics were similarly calculated for each of the classical benchmark models.

### Sliding window positional analysis

To investigate the importance of each amino acid within the 9-mer peptide sequence, we implemented a sliding window approach. Each model was designed to take four specific peptide positions as input features. To determine which positions were most informative for either binding or immunogenicity classification, we systematically evaluated all possible combinations out of the nine total (i.e. 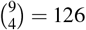 combinations). For each set of unique positions, a separate model was trained and evaluated. All models used identical training and testing splits for their respective classification. This allowed us to analyze the contribution of different regions of the peptide and identify which positions were most predictive of the respective biological outcome.

### Survival analysis

When examining if our model was able to stratify patients based on their predicted immunogenic peptide burden, we utilized a large scale clinical trial of immunotherapy treated lung cancer patients^29^. From these patients whole exome sequencing derived somatic mutations, we identified 214,981 peptides with the potential to form immunogenic antigens across 111 patients with HLA-A02:01. The data set was preprocessed and encoded as previously described, leaving 209,889 unique neoantigen peptides that were fully processed (i.e. did not contain missing peptide information such as a ‘*’ or ‘X’). To develop our Q-CHIPP model we combined two pretrained models. The first model, trained on 150 peptides from the IEDB dataset, used positions 2, 4, 5, and 9 to predict MHC binding. The second model, trained on 300 confirmed MHC-binding peptides, used positions 3, 5, 6, and 7 to predict immunogenicity. To aggregate their outputs, we applied a bit-wise AND operation to both sets of predicted labels; therefore, Q-CHIPP predicts positive only when both individual models predict a positive outcome. After all peptides were analyzed and classified, the predictions were grouped by patient and the total number of immunogenic peptides were calculated for each patient. This process was repeated for both classical CNN models (CHIPP-limited and CHIPP-large) as well as for the random forest combination model. For each, we loaded in the respective pretrained models, tested each on the^29^ dataset to get the predicted labels, and then aggregated the two individual models. In addition to these models, for each peptide we used NetMHCpan’s BA RANK to classify each peptide as a likely binder using a threshold of 2.0.

To evaluate the association between survival outcomes and immunogenicity-related metrics, we conducted a Kaplan-Meier survival analysis using `lifelines` Python package. We focused on overall survival (OS) as the primary clinical endpoint, using the time duration the patient survived from the start point until death or last follow-up (OS_months). For each metric under investigation (NetMHC, QCNN, CCNN limited, and CCNN large), we stratified patients into two groups: *High* and *Low* based on the median value of the respective metric. Specifically, patients with a value greater than or equal to the median were assigned to the High group and those below to the Low group. Survival curves for the High and Low group were calculated using the Kaplan-Meier estimator. Differences between groups were assessed using the log-rank test, with the resulting p-values reported for each comparison. Survival plots were then generated to visualize the survival probabilities over time along with risk tables showing the number of individuals at risk at different time points.

### Enrichment Comparison

To compare amino acid enrichment between positive and negative predictions for each model (Q-CHIPP, CHIPP-limited, CHIPP-large), predicted labels were loaded from their respective outputs. Each 9-mer peptide was one-hot encoded into a 1×180 binary vector, representing the presence of each amino acid at each position. To compare the enrichment of positive predictions against negative predictions, (Fig. 6 F-H) peptides were subset to divided into two groups: peptides that were predicted as immunogenic positive and those that were predicted to be negative. For both the positive and negative group, we summed the one-hot encoded amino acid values for the respective group. These summed values were normalized by the total number of samples for each group. We then compute the per-feature ratio of the positive values to each of the negative values for each model. To prevent division by zero, we added a small constant (*ε* = 1e-6) to both numerator and denominator. Each feature was decomposed into its corresponding amino acid and position and we restricted analysis to positions 2, 3, 4, 5, 6, 7, and 9 as each model’s input was subset to only these positions (2, 4, 5 and 9 and 3, 5, 6, and 7). We then applied a log-scale normalization using base 2 with a small constant to avoid log(0). Values were then clipped to the range [-3, 3]. Each heatmap was then developed using `seaborn.heatmap` library for each model.

### Benchmark models

Two classical CNNs implemented through `tensorflow` (v2.16.1) were used to compare performance against the QCNN. The first model was engineered to mimic the limited number of parameters seen within the quantum CNN design. Additionally, we constructed a unrestricted CNN model that gave us a benchmark on the full performance on the classical side. All models were initialized with a specified seed (49) for both `numpy` and `tensorflow`. Each of the train and test datasets used for the classical models were identical to the splits used in the quantum models. Furthermore, the training and testing data was loaded in and processed in the same way as previously described.

To construct the limited classical CNN model we use Keras’ `Sequential` API. The model was designed to adapt its number of parameters based on the input sequence length. Specifically, the model dynamically chooses the number of convolutional layers (`Conv1D`) and number of filters for each layer based on the input size. The input size consists of the number of bits encoded and ranges for the number of features encoded. For instance, for inputs of length ≤12m the model uses two convolutional layers with 2 filter each; for lengths *>* 12 and ≤24, it uses three layers with 4 filters each; and for longer sequences it gradually scales up to 4, 8, and 8 filters. All convolutional layers use a kernel size of 3, ‘same’ padding, ReLU activation, and L2 regularization (0.0005). Each convolutional layer is followed by a max-pooling (`MaxPooling1D`) with a pool size of 2. The final convolutional output is passed through a `GlobalAveragePooling1D` layer to flatten the dimension while minimizing overfitting. A single sigmoid-activated `Dense` layer is then used to produce the final binary output. The model was then compiled using the Adam optimizer with binary cross entropy loss.

The unrestricted, larger CNN model was designed with three one-dimensional convolutional layers (`Conv1D`) using 32, 54 and 128 filters, respectively. Each layer had a kernel size of 3, used ReLU activations, and was implemented using L2 regularization (0.0005). The initial two convolutional layers were followed by a `MaxPooling1D` layer with a pooling size of 2. The final convolutional layer was followed by a `GlobalAveragePooling1D` layer to reduce overfitting while preserving spacial information. Following the convolutional layers, two fully connected `Dense` layers were implemented with 128 and 64 units, respectively. Both layers used ReLU activations and dropout regularization (rate=0.3). A final dense layer with sigmoid activation was then used to produce the binary classification output. The model was then compiled using the Adam optimizer with binary cross entropy loss.

We additionally benchmarked the QCNN model against a standard random forest model. This model was chosen due to its simplistic architecture and is utilized as a standard benchmark in drug discovery applications. We initially pre-processed each peptide to contain the peptide sequence and HLA (which is constant as we are strictly analyzing A0201 HLA types). For each peptide, we converted each character to its ASCII value and then selected the desired peptide positions per sequence. These numeric features were one-hot encoded and combined to form the input to the model. We used `RandomForestClassifier` from `sklearn` with 300 trees and default hyper-parameters. The model was trained on the processed training data and evaluated on the test set. To assess the variability in the random forest classifier, we repeated training and evaluation 100 times. The final performance was measured using Area Under the ROC Curve (AUC) and F1 for each run and then averaging all 100 iterations.

## Data and Code availability

Data sets used throughout this paper include those from IEDB, our previous published work on neoantigens, and public data from EGA datasets EGAD00001011302, EGAS00001007509, EGAS00001007508, and EGAS00001003892.

## Acknowledgments

The authors acknowledge the support from the Cleveland Clinic - IBM Discovery Accelerator, and the use of IBM Quantum System One at Cleveland Clinic. Part of this material is based upon work supported by the National Science Foundation Grant No. DBI-1548297

## Author contributions

Conceptualization, T.J.A., F.U., S.C., L.P., J.K.W., T.A.C, R.P., K.R., P.P.; methodology,F.U., R.P., K.R., P.B.P., T.G., M.T., J.S., J.K.W., T.J.A., V.R.; investigation, R.P., T.J.A., P.B.P., F.U., K.R., V.R., T.G., M.T., J.S., J.K.W., S.C., L.P.; writing – original draft, T.J.A, R.P., P.B.P., K.R., F.U., S.C.; Writing – review & editing, T.J.A, R.P., P.B.P., K.R., F.U., L.P., S.C., T.A.C., T.G., M.T., J.S., J.K.W., V.R.; funding acquisition, T.J.A., S.C.; supervision, T.J.A., F.U., K.R., S.C., L.P., T.A.C.; All authors critically discussed and revised the manuscript for important intellectual content.

---

**Extended Data Fig 1.**
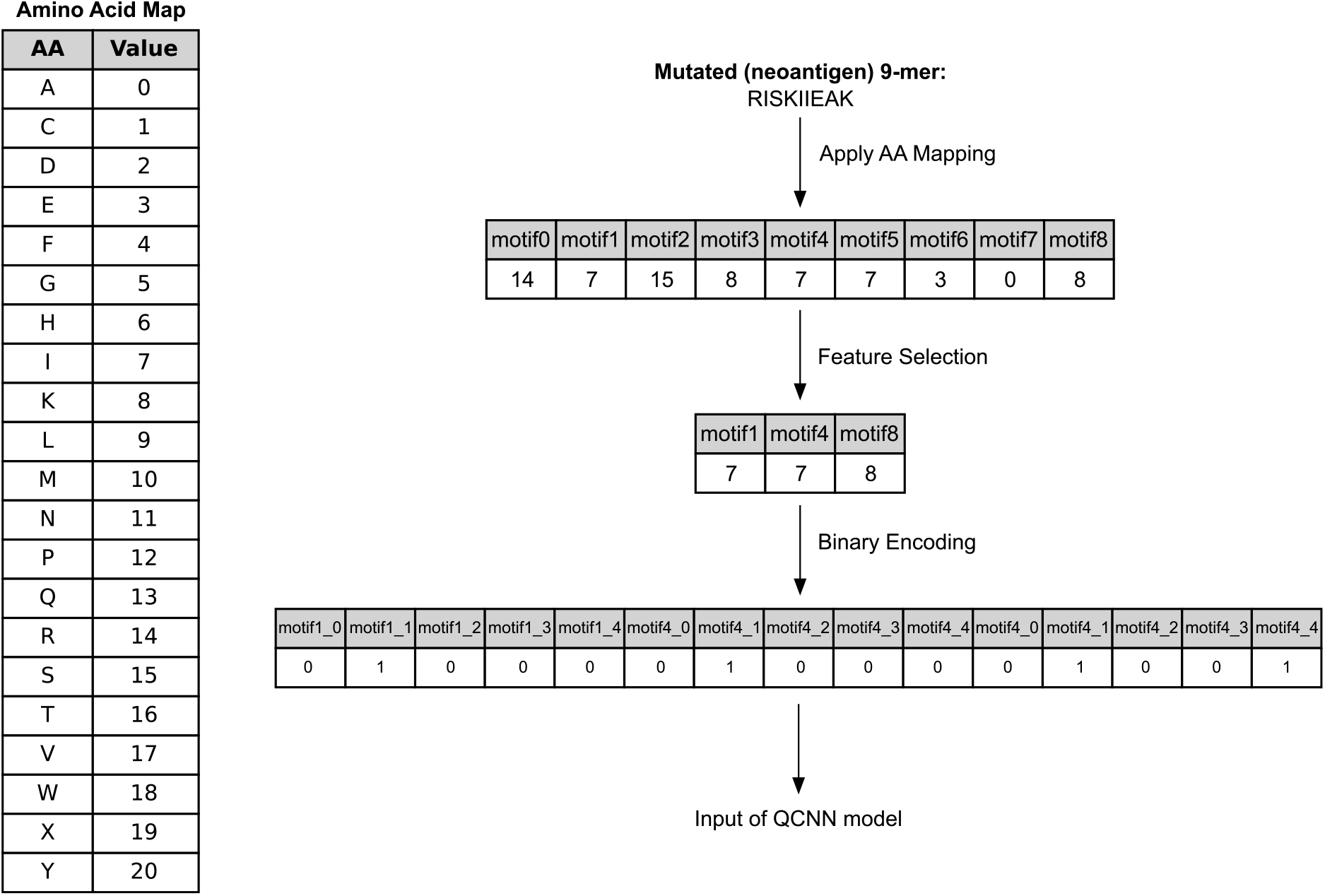
Peptide encoding pipeline for QCNN model. Schematic of the data preprocessing steps used to convert a given mutated peptide (9-mer sequence of amino acids) to its respective input for the QCNN model. For each peptide, we translate each amino to its respective integer using the amino acid map. This sequence is subset to include only features used for a given model (peptide positions 2, 5, and 9 which are 1, 4 and 8 when zero indexed). Binary encoding is then applied to the dataset to map it to a 5 bit vector for each amino acid. This sequence is then passed as input for the model.

**Extended Data Fig 2.**
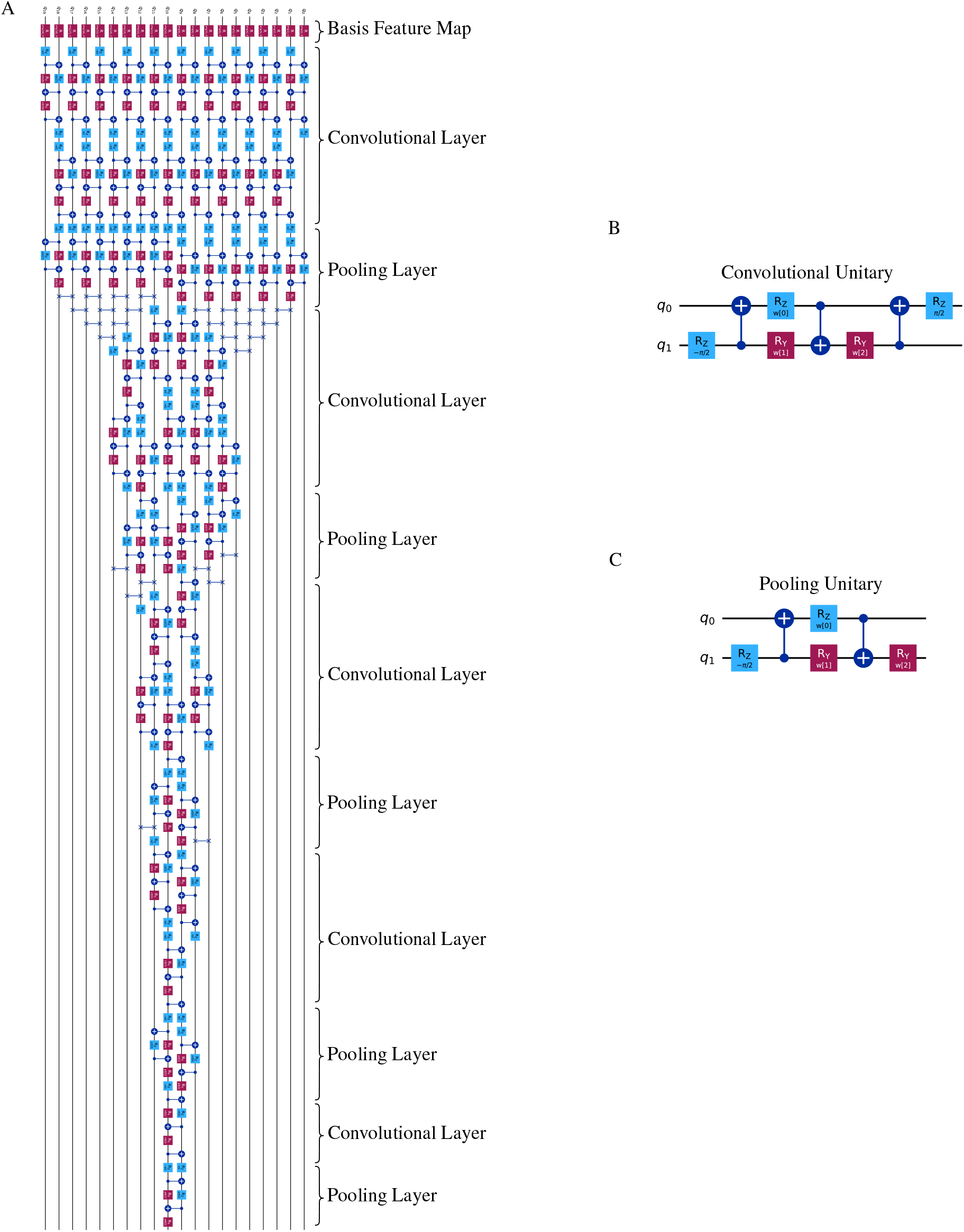
Architecture of the 20-qubit QCNN. A) Schematic of the full QCNN used in the 20-qubit experiment. The model consists of an initial feature mapping followed by a series of convolutional and pooling unitaries. B) Detailed circuit of each convolutional unitary. C) Detailed circuit of each pooling unitary.

**Extended Data Fig 3.**
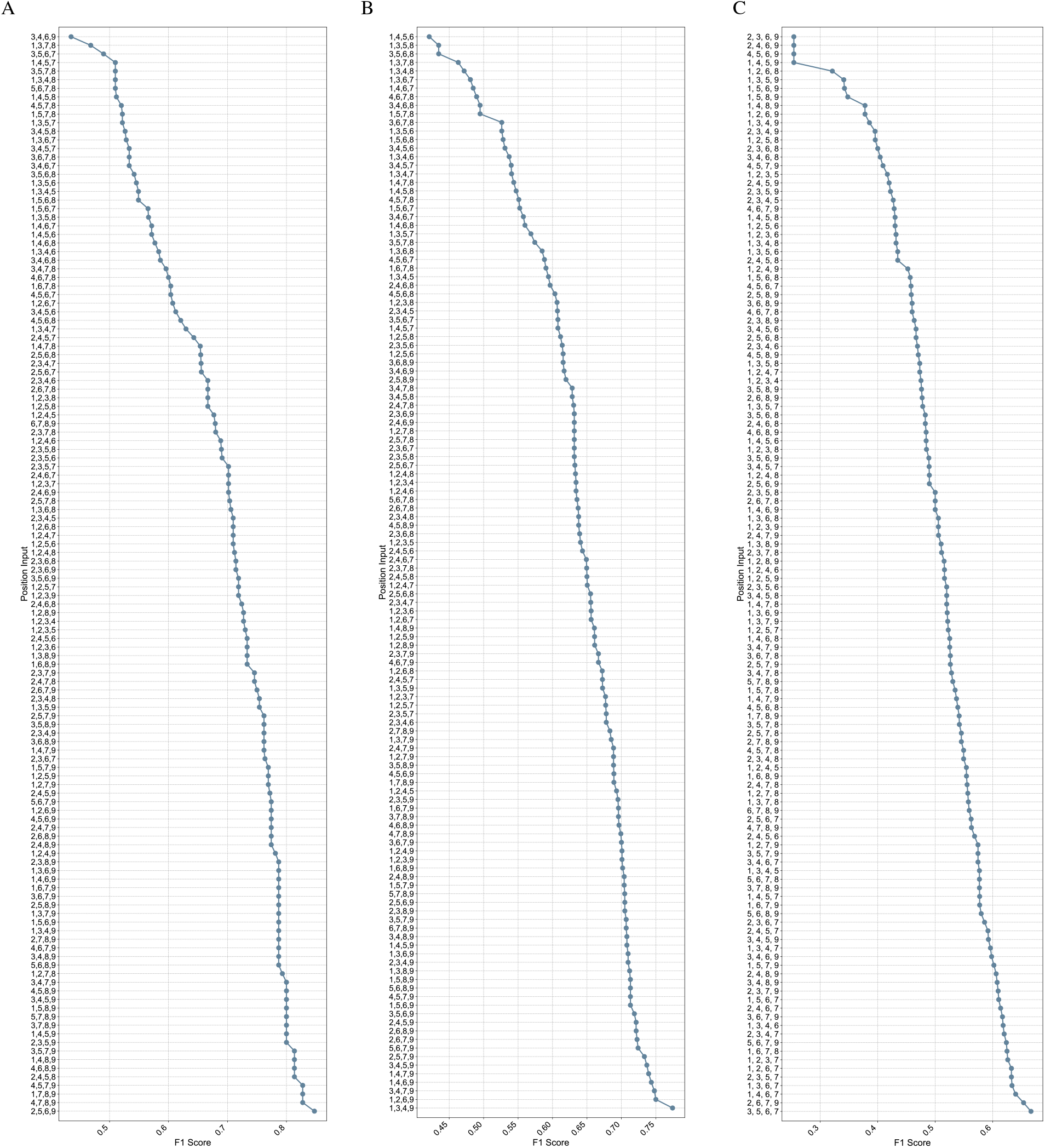
QCNN performance across input feature sets for binding and immunogenicity prediction. F1 performance of the QCNN model given unique a set of 4 positions as input. Three listed models include A) binding predictions, B) immunogenicity predictions given even split of both binders and non-binders, and C) immunogenicity predictions given subsample of confirmed MHC binders. All three experiments were trained and tested using the noiseless statevector simulator with QRAC 2:1 feature mapping.

**Extended Data Fig 4.**
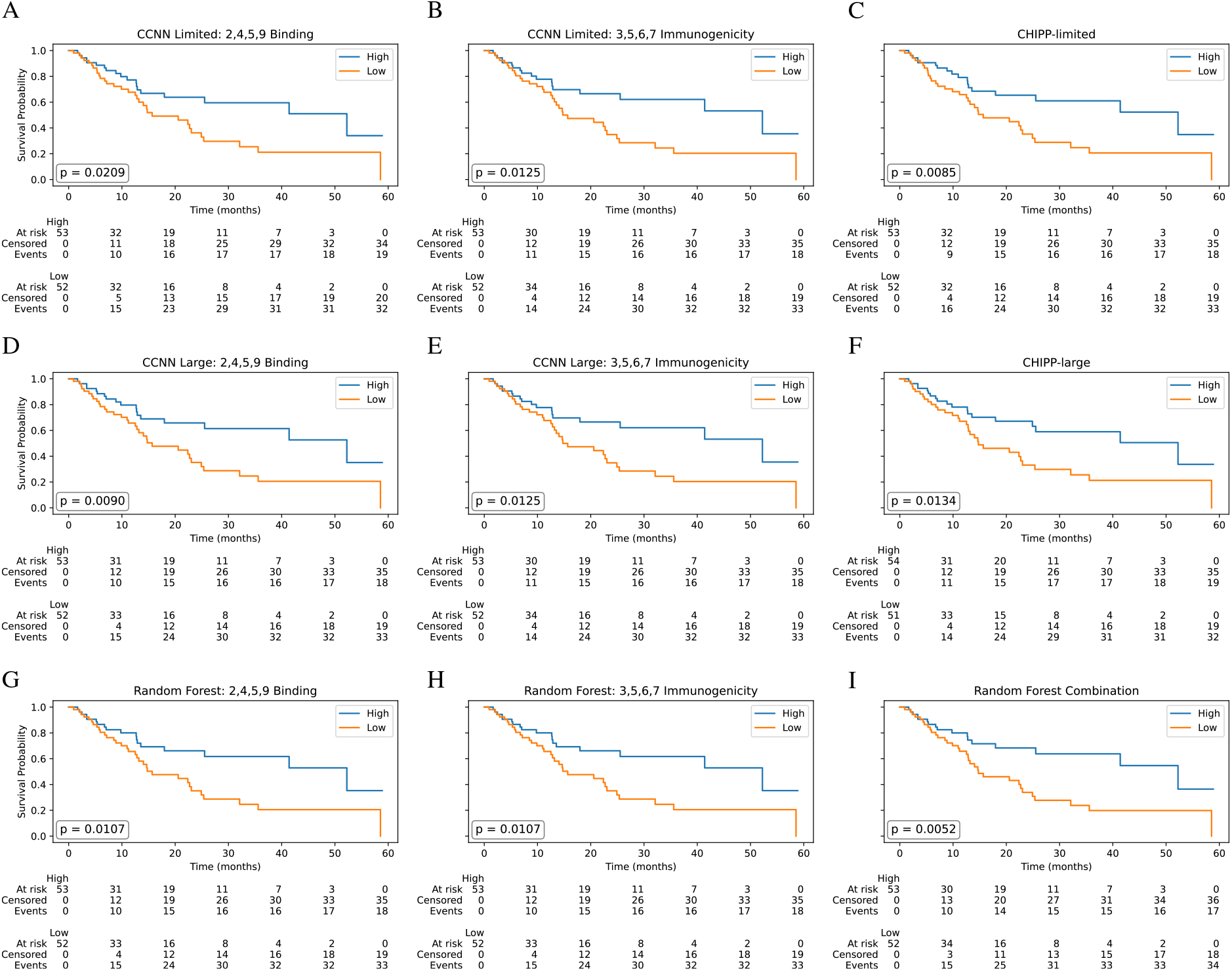
Patient survival estimations using classical methods. Ability of each model to separate patents based on overall survival. For each individual benchmark, survival was evaluated using a model trained to predict binding using an IEDB dataset with positions 2, 4, 5, and 9. Additionally used was a model trained to predict immunogenicity from those peptides confirmed to bind to MHC from IEDB dataset using positions 3, 5, 6, and 7. Finally both models were aggregated to create a combination of both models. Each model was trained on identical data splits. Depicted is the binding, immunogenicity, and combinatory model for the CCNN limited model (A-C), CCNN large model (D-F), and random forest (G-I).

